# Dependence on a variable residue limits the breadth of an HIV MPER neutralizing antibody, despite convergent evolution with broadly neutralizing antibodies

**DOI:** 10.1101/2022.03.18.484856

**Authors:** Cathrine Scheepers, Prudence Kgagudi, Nonkululeko Mzindle, Elin S. Gray, Thandeka Moyo-Gwete, Bronwen E. Lambson, Brent Oosthuysen, Batsirai Mabvakure, Nigel J. Garrett, Salim S. Abdool Karim, Lynn Morris, Penny L. Moore

**Affiliations:** Centre for HIV and STIs, National Institute for Communicable Diseases of the National Health Laboratory Service, Johannesburg, 2131, South Africa; SA MRC Antibody Immunity Research Unit, School of Pathology, University of the Witwatersrand, Johannesburg, 2050, South Africa; Centre for the AIDS Programme of Research in South Africa (CAPRISA), KwaZulu-Natal, 4013, South Africa; Department of Epidemiology, Columbia University, New York City, NY, 10032, USA; Institute of Infectious Disease and Molecular Medicine, University of Cape Town, Cape Town, 7701, South Africa

## Abstract

Broadly neutralizing antibodies (bNAbs) that target the membrane-proximal external region (MPER) of HIV gp41 envelope, such as 4E10 and VRC42.01, can neutralize >95% of viruses. These two MPER-directed monoclonal antibodies share germline antibody genes (*IGHV1-69* and *IGKV3-20)* and form a bNAb epitope class. Furthermore, convergent evolution within these two lineages towards a ^111.2^GW^111.3^ motif in the CDRH3 is known to enhance neutralization potency. We have previously isolated an MPER neutralizing antibody, CAP206-CH12, that uses these same germline heavy and light chain genes but lacks breadth (neutralizing only 6% of heterologous viruses). Longitudinal sequencing of the CAP206-CH12 lineage over three years revealed similar convergent evolution towards ^111.2^GW^111.3^ among some lineage members. Mutagenesis of CAP206-CH12 from ^111.2^GL^111.3^ to ^111.2^GW^111.3^ modestly improved neutralization breadth (by 9%) and potency (3-fold), but did not reach the levels of breadth and potency of VRC42.01 and 4E10. To explore the lack of potency/breadth, viral mutagenesis was performed to map the CAP206-CH12 epitope. This indicated that CAP206-CH12 is dependent on D^674^, a highly variable residue at the solvent-exposed elbow of MPER. In contrast, VRC42.01 and 4E10 were dependent on highly conserved residues (W^672^, F^673^, T^676^, and W^680^) facing the hydrophobic patch of the MPER. Therefore, while CAP206-CH12, VRC42.01, and 4E10 share germline genes and show some evidence of convergent evolution, their dependence on different amino acids, which impacts orientation of binding to the MPER, result in differences in breadth and potency. These data have implications for the design of HIV vaccines directed at the MPER epitope.

**Author Summary:** Germline-targeting immunogens are a promising HIV vaccine design strategy. This approach is reliant on the identification of broadly neutralizing antibody (bNAb) classes, which use the same germline antibody genes to target the same viral epitopes. Here, we compare three HIV Envelope MPER-directed antibodies (4E10, VRC42.01 and CAP206-CH12) that despite having shared antibody genes, show distinct neutralization profiles. We show that CAP206-CH12 targets a single highly variable residue in the MPER, which results in low neutralization breadth. In contrast, the 4E10 and VRC42.01 mAbs are dependent on highly conserved residues in the MPER, resulting in exceptional neutralization breadth. Our data suggest that while shared germline genes within bNAb epitope classes are required, in some cases these are not sufficient to produce neutralization breadth, and MPER immunogens will need to trigger responses to conserved sites.

## INTRODUCTION

The pursuit of an effective vaccine against HIV is an ongoing priority. It is generally accepted that an effective vaccine will require the elicitation of broadly neutralizing antibodies (bNAbs), capable of neutralizing multiple subtypes of HIV [1]. Recent results from the antibody-mediated prevention (AMP) trials demonstrated that passive infusion of the VRC01 bNAb prevented infection by viruses sensitive to this antibody re-invigorating the search for bNAb-inducing vaccines [2]. However, eliciting bNAbs by vaccination has proven to be challenging because they develop only in ~25% of infected donors even after many years and tend to have unusual features such as high levels of somatic hypermutation (SHM), long heavy chain or short light chain third complementarity determining regions (CDRH3s/CDRL3s) [3]. The identification of bNAb classes, which share common germline antibody genes and target the same region on the HIV envelope, has resulted in several germline-targeting vaccine strategies that aim to trigger unmutated common ancestors (UCAs) of bNAbs [4–9]. Studies defining bNAb/virus co-evolution during HIV infection have been invaluable in revealing the characteristics of early precursors/unmutated common ancestors (UCA), antibody intermediates, and the viral variants that engage and drive these lineages [10–19].

The 4E10 bNAb class, including 4E10 and VRC42.01, are amongst the broadest antibodies described to date (>95% of multi-subtype virus panels) [15,20]. These bNAbs target the membrane proximal external region (MPER) of the HIV-1 gp41 envelope glycoprotein, sharing some dependence on the tryptophan (W) at residue 680 [15,20]. 4E10 and VRC42.01 use the same heavy and light chain germline genes: *IGHV1-69* and *IGKV3-20*, with modest SHM (heavy chains: 8.3 and 11.5%, respectively; light chains: 5.3 and 5.7%, respectively). In addition to shared germline gene usage, 4E10 and VRC42.01 show convergent evolution within the CDHR1 and CDRH3. Within the CDRH1, the ^25^SGGSFS^30^ motif that is crucial for binding, is largely encoded in all germline *IGHV1-69* alleles, with only the S^28^ being mutated within 4E10 and VRC42.01, since all *IGHV1-69* germline alleles contain a T^28^ [20]; (www.imgt.org). Within the CDRH3, VRC42.01 and 4E10 mutated towards a ^111.2^GW^111.3^ motif (IMGT numbering), with 4E10 having a double ^111^GWGW^111.3^ motif, which is crucial for its neutralization [15,20].

We have previously reported the isolation of mAbs from donor CAP206 who developed broadly neutralizing plasma responses to the MPER [21]. CAP206-CH12 was isolated at 120 weeks post-infection (wpi) and an early intermediate of the same lineage, CAP206-CH12.2, from 17 wpi [22,23]. Like 4E10 and VRC42.01, the CAP206-CH12 lineage makes use of the *IGHV1-69* and *IGKV3-20* germline genes and targets the MPER [22]. However, despite identical germline variable gene usage and an overlapping epitope, CAP206-CH12 shows limited neutralization breadth (~6% of heterologous viruses) compared to the two bNAbs [15]. Here, we use longitudinal antibody deep sequencing of the CAP206-CH12 lineage over three years to understand the lack of breadth and potency in this lineage compared to 4E10 and VRC42.01.

## RESULTS

We have previously described the kinetics of the MPER response in donor CAP206, using an HIV-2/HIV-1 MPER chimera involving an HIV-2 backbone (7312A) with an HIV-1 MPER region, showing that plasma neutralization against the MPER was detected from 25 wpi and seen throughout infection [24] (**Fig 1A**). To study the evolution of the CAP206-CH12 lineage, we carried out antibody next-generation sequencing using gene-specific heavy and light chain primers (*IGHV1-69* and *IGKV3-20*) from seven-time points (10, 33, 56, 94, 120, 185, and 198 wpi) during infection (**Fig 1A**).

**Fig 1.**
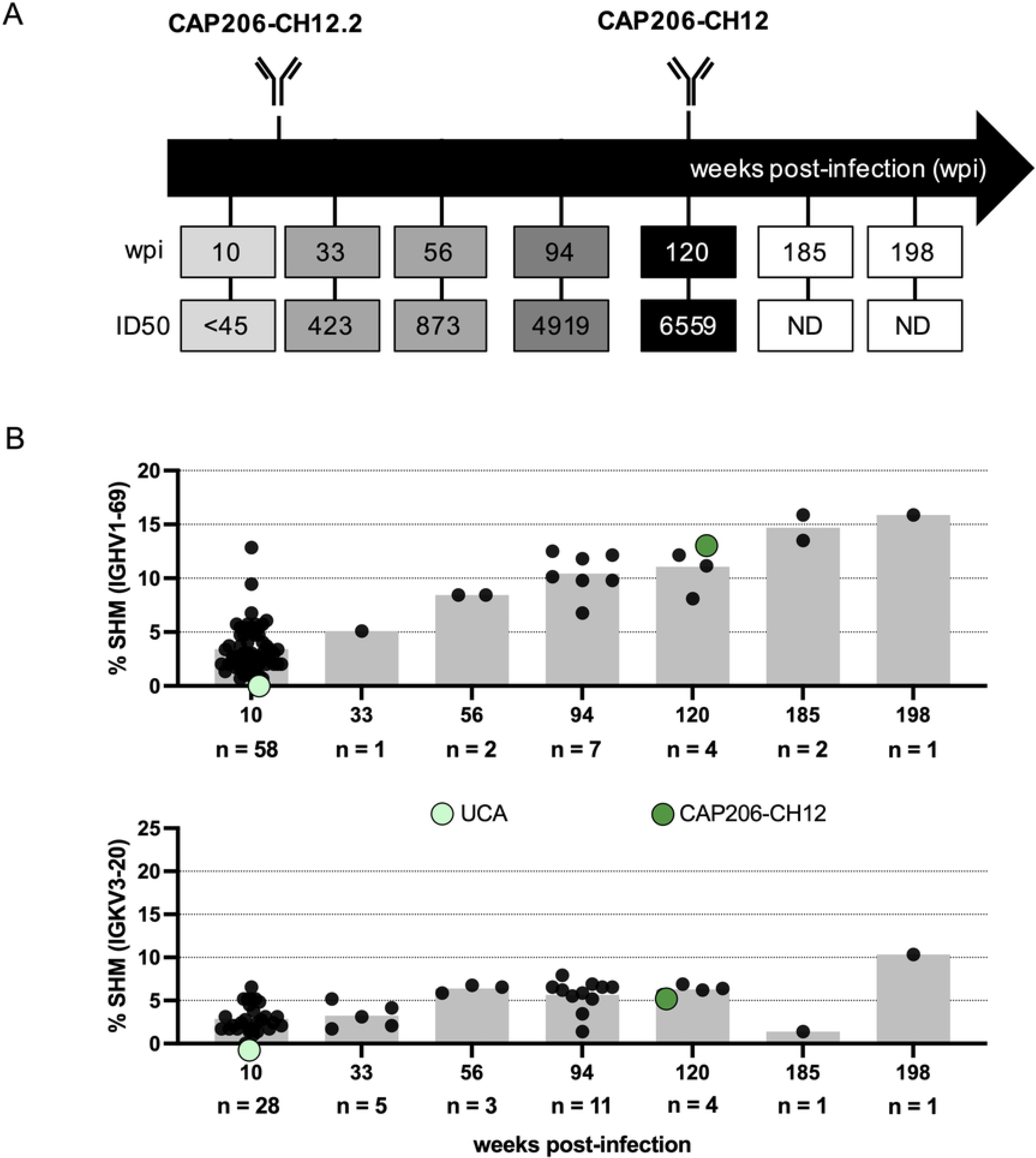
CAP206-CH12 lineage is detected throughout infection. **(A)** Time-points at which longitudinal antibody next-generation sequencing (Illumina-MiSeq) were conducted and CAP206-CH12.2 and CAP206-CH12 mAbs were isolated. ID_50_ plasma neutralization titres of C1C, indicative of an MPER response, are shown in grey scale. **(B)** Percentage of heavy and light chain somatic hypermutation (SHM) obtained from clonally related sequences from each time-point, compared to their germline *IGHV1-69* and *IGKV3-20* genes. The grey bars at each time point represent the average percentage SHM. Each dot represents a single heavy or light chain clonally related sequence identified by time-point. The total number of sequences per time-point are given on the x-axis. The levels of SHM for the UCA and CAP206-CH12 are shown as green dots on the graph.

Clonally related sequences were detected at all time-points sequenced for both heavy (n = 75) and light chains (n = 53). The majority of the clonally related reads were obtained at 10 wpi for both heavy and light chains, with fewer sequences detected at later time-points (**Fig 1B**). Heavy and light chain sequences with no SHM, representing the UCA (light green) of this lineage, were detected at 10 wpi. Somatic hypermutation increased with the duration of infection, with the heavy chain being more mutated than the light chain, reaching an average of 15.8% and 10.3%, respectively (**Fig 1B**).

### Neutralization is largely mediated by CAP206-CH12 heavy chain

We identified two light chain sequences from the same time-point (120wpi, K1) as CAP206-CH12 or later (198wpi, K2) that were more mutated than CAP206-CH12 (**Fig 2A and B**). In an attempt to improve potency of CAP206-CH12 we paired K1 (6.4% SHM, light purple) and K2 (10.3% SHM, dark purple) with CAP206-CH12 heavy chain.Neutralization of eight heterologous subtype B and C viruses (CAP85, COT6, Du156, Du422, QHO692, TRO.11, ZM135, and ZM197) was compared to the CAP206-CH12 wild-type (**Fig 2C**). For the majority of these viruses, very little or no difference was seen between the CAP206-CH12 heavy chain paired with different light chains. For two of the viruses (Du156 and TRO.11), the most mature light chain K2 showed reduced neutralization compared to CAP206-CH12. This indicates that increased SHM within the light chain doesn’t improve the neutralization of CAP206-CH12.

**Fig 2.**
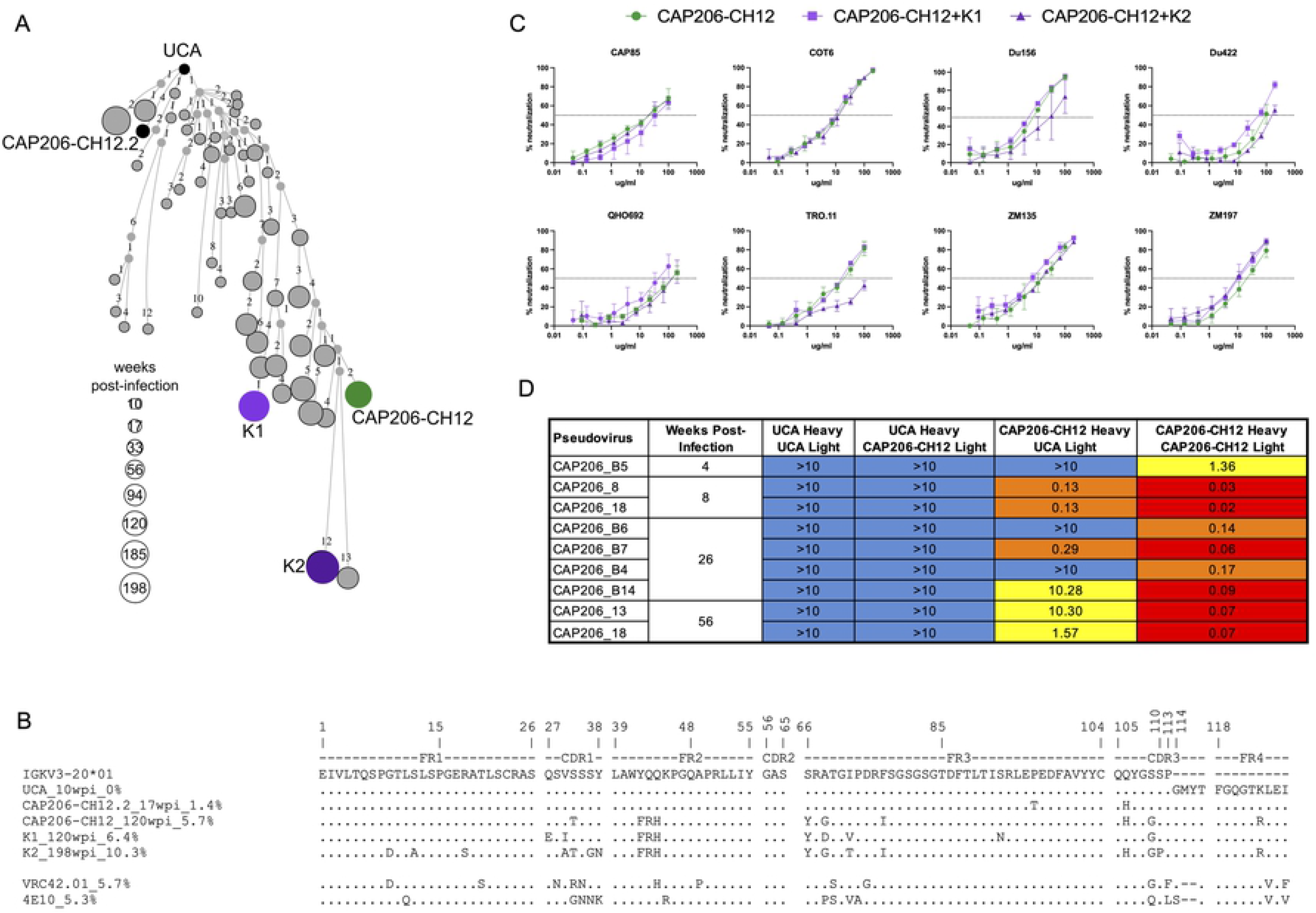
CAP206-CH12 lineage light chain plays a minor role in neutralization. **(A**) CAP206-CH12 lineage reconstruction tree of light chain clonally related sequences. Small nodes present intermediate sequences inferred by Change-O, part of the immcantation portal, while larger circles represent clonally related sequences observed in the NGS data, sized according to weeks post-infection. The UCA, CAP206-CH12.2, CAP206-CH12 as well as two additional light chains (K1 and K2), used for further analysis, are labeled and shown as black or coloured circles on the tree. **(B)** Alignment of light chains of the UCA, CAP206-CH12.2, CAP206-CH12, K1, K2, VRC42.01, and 4E10 against their shared *IGKV3-20* germline gene. Sequences are numbered according to the IMGT numbering system (https://www.imgt.org). **(C)** Neutralization curves of CAP206-CH12 with its natively paired heavy and light chains (green) in comparison to CAP206-CH12 heavy chain paired with either K1 (light purple) or K2 (dark purple) against eight heterologous viruses. Shown are the average neutralization titres (IC_50_ in μg/mL) with the error bars representing standard deviations. **(D)** Neutralization of autologous viruses from the first year of infection by the UCA, CAP206-CH12, and UCA/CAP206-CH12 heavy and light chain chimeras. The table represents IC_50_ titres and are colored according to titre where blue represents the inability to neutralize the virus, yellow titres between 1 – 10 μg/mL, orange between 0.1 – 0.9 μg/mL, and red between 0.01 – 0.09 μg/mL.

To confirm that the light chain played a minor role in CAP206-CH12 neutralization, chimeras of the UCA and CAP206-CH12 heavy and light chains were tested against autologous viruses isolated within the first year of infection, and compared to the wild-type UCA and CAP206-CH12 (**Fig 2D**). Both the wild-type UCA and the UCA heavy chain paired with the CAP206-CH12 light chain showed no neutralization against any of the autologous viruses. In contrast, the CAP206-CH12 heavy chain paired with the UCA light chain showed some neutralization of autologous viruses, though with less potency than the CAP206-CH12 wild-type antibody. This indicates that the light chain plays a minor role in CAP206-CH12 neutralization, with the majority of neutralization potency mediated by the heavy chain.

### “^111.2^GW^111.3^” motif in the CDRH3 improves neutralization breadth and potency in CAP206-CH12 lineage

We, therefore, focused on the heavy chain maturation and specifically the CDRH3 as both VRC42 and 4E10 lineages showed convergent evolution towards a ^111.2^GW^111.3^ motif, which is crucial for neutralization [15,20]. We observed convergent evolution within the CAP206-CH12 lineage towards the same motif, with the sequences seen in the later time-points having mutated away from the ML motif seen in the UCA (grey) to either a ^111.2^GL^111.3^ (orange) or ^111.2^GW^111.3^ (red) (**Fig 3A** and **B**). Notably, none of the CAP206-CH12 lineage members contained a double ^111^GWGW^111.3^ motif as seen in 4E10 or a ^111^GAGW^111.3^ as seen in VRC42.01 (motif highlighted in grey, **Fig 3B**). In addition to the CDRH3 ^111.2^GW^111.3^ motif, we observed maturation within other parts of the heavy chain, particularly within the CDRH1, FR2, and CDRH2, with CAP206-CH12.2 sharing the ^25^SGGSFS^30^ motif, crucial for binding for VRC42.01 and 4E10, though the majority of the CAP206-CH12 intermediates maintained the germline-encoded T^28^ (**Fig 3B**).

**Fig 3.**
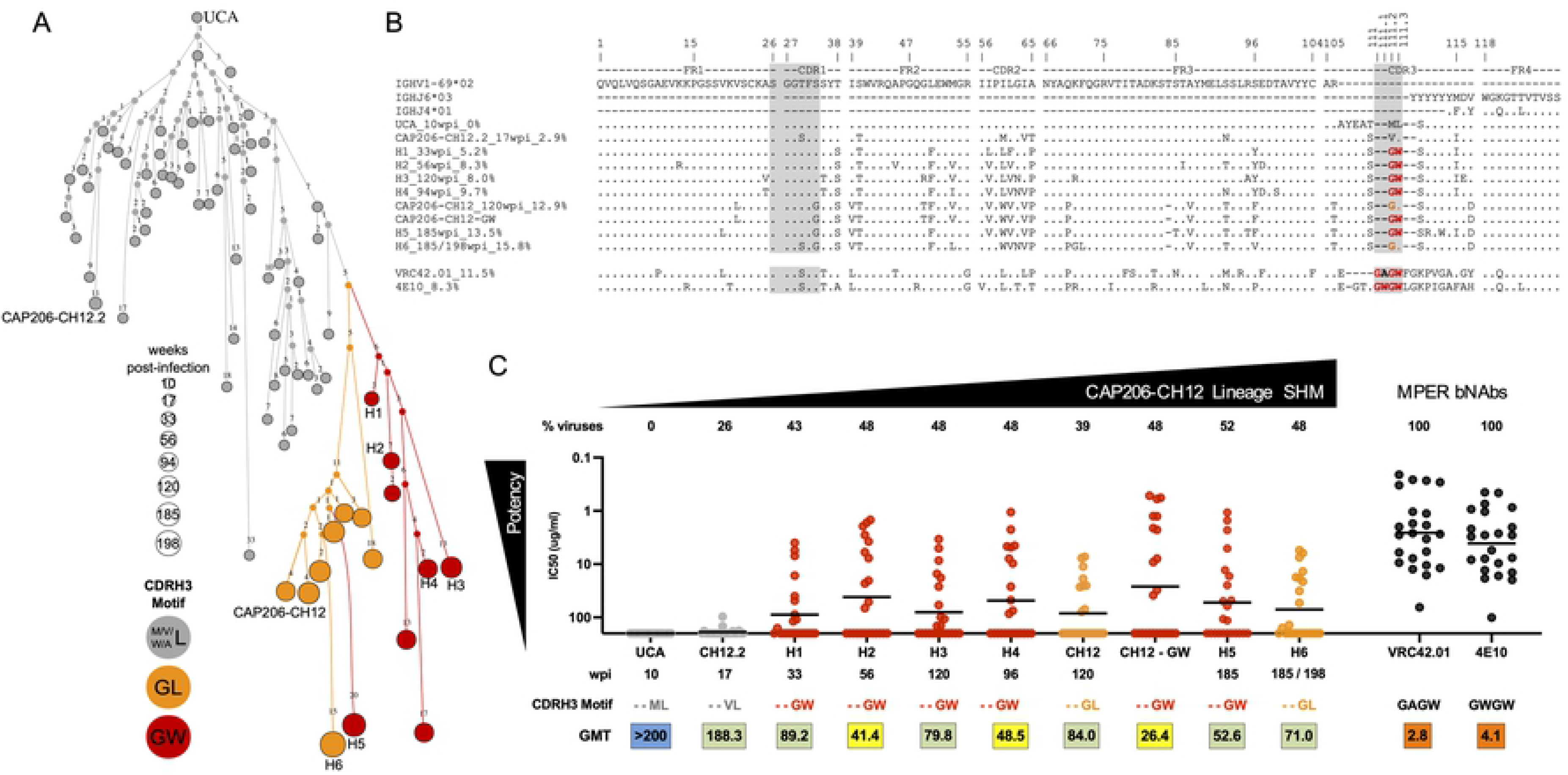
“^111.2^GW^111.3^” motif increases neutralization potency. **(A)** Reconstruction tree of the CAP206-CH12 clonally related heavy chain sequences colored according to CDRH3 motif at positions 111.2 and 111.3. Those having a methionine, valine, tryptophan, or alanine (M/V/W/A^111.2^) and leucine (L^111.3^) are shown in grey, those with a glycine and leucine (^111.2^GL^111.3^) in orange, and those with a glycine and tryptophan (^111.2^GW^111.3^) in red. Small nodes present intermediate sequences inferred by Change-O, part of the immcantation portal, and clonally related sequences observed in the NGS data are represented by larger circles, sized according to time-point. Sequences selected for testing (UCA, H1 – H6, CAP206-CH12.2, and CAP206-CH12) are labeled. **(B)** Heavy chain sequences of the UCA, CAP206-CH12.2, H1 – H6, CAP206-CH12, VRC42.01, and 4E10 compared to the shared germline *IGHV1-69* and *IGHJ6* (*IGHJ4* used by VRC42.01 and 4E10). CDRH1 and CDRH3 regions important for 4E10 and VRC42.01 binding are highlighted in grey. Within the CDRH3, amino acids are colored according to the motif as shown for (C). Sequences are numbered according to the IMGT numbering system (https://www.imgt.org). **(C)** Neutralization of CAP206-CH12 lineage mAbs (colored according to the ^111.2^GL^111.3^ or ^111.2^GW^111.3^ motif as seen in panels A and B) and MPER bNAbs (VRC42.01 and 4E10) against a panel of 23 viruses (neutralized by CAP206 plasma). The black line for each mAb represents the geometric mean titre. The percentage of viruses (out of a panel of 23 viruses that were neutralized by CAP206 donor plasma [21]) neutralized are shown above each mAb. The time-point at which each heavy chain sequence was observed or mAb was isolated, CDRH3 motif, and geometric mean titre are given below each mAb.

In order to test the influence of the ^111.2^GW^111.3^ motif within the CAP206-CH12 lineage, we selected six intermediate heavy chain sequences (H1 - H6), including those with either ^111.2^GL^111.3^ (orange, **Fig 3**) or ^111.2^GW^111.3^ motifs (red). Since CAP206-CH12 has a ^111.2^GL^111.3^ motif we also mutated the L^111.3^ to a W^111.3^ (referred to as CAP206-CH12-GW). We tested the ability of these antibodies including the UCA, CAP206-CH12.2, and wild-type CAP206-CH12 to neutralize a panel of 23 viruses that we have previously shown to be sensitive to CAP206 plasma [24], and compared these to VRC42.01 and 4E10 (**Fig 3C**). The UCA of the CAP206-CH12 lineage was unable to neutralize any of the heterologous viruses, while the early intermediate CAP206-CH12.2 showed limited neutralization (6/23 viruses, 26% breadth). As with many antibody lineages, increased SHM resulted in increased neutralization breadth [10–19]. The mutated CAP206-CH12-GW showed statistically significantly greater breadth and potency (p=0.001) compared to the wild-type CAP206-CH12 (with the ^111.2^GL^111.3^ motif) (**Fig 3C**). Furthermore, antibodies with lower SHM compared to CAP206-CH12 but having the ^111.2^GW^111.3^ motif (H2 - H4) showed greater breadth and potency than CAP206-CH12. Similarly, H6, the most mature mAb (15.8% SHM) but containing the ^111.2^GL^111.3^ motif, showed weaker neutralization breadth and potency compared to a less mature mAb (H5, 13.5% SHM) present at the same time-point but with the ^111.2^GW^111.3^ motif. This data suggests that the ^111.2^GW^111.3^ motif improves neutralization potency in the CAP206-CH12 lineage, as with VRC42.01 and 4E10 [15].

Despite this, none of the lineage members showed the same degree of neutralization breadth or potency as VRC42.01 or 4E10 (GMT: 2.8 and 4.1 ug/mL, respectively). Furthermore, the CAP206-CH12 lineage reached a plateau in neutralization at 52% with most mAbs only able to neutralize 48% of viruses sensitive to the CAP206 plasma, from whom this lineage was derived. This suggests additional antibody lineages exist within this donor that account for the remaining plasma neutralization breadth.

### Multiple MPER lineages and specificities detected in donor CAP206 plasma

We performed a longitudinal assessment of CAP206 plasma neutralization of all 23 heterologous viruses neutralized by this donor and partitioned the viruses into those sensitive (green) and those resistant (black) to the CAP206-CH12 lineage (**Fig 4A**). Neutralization of lineage-sensitive viruses was detected from 15 wpi and persisted until 159 wpi. This is consistent with the timing of the CAP206-CH12 lineage, with the UCA detected from 10 wpi, isolation of mAbs CAP206-CH12.2, and CAP206-CH12 at 17 and 120 wpi respectively, and the most mature mAb detected at 198 wpi. Furthermore, CAP206-CH12.2 is capable of neutralizing UG024 and 92RW, which are the first viruses neutralized by the plasma from 15 wpi. The emergence of the ^111.2^GW^111.3^ motif at 33 wpi (detected in H1 - H4 and CAP206-CH12) enabled the lineage to neutralize additional heterologous viruses, with all the lineage-sensitive viruses neutralized by 80 wpi. Interestingly, with the exception of ZM109, neutralization of all lineage-resistant viruses (black) developed after 80 wpi, suggesting another lineage or specificity emerged at that time-point.

**Fig 4.**
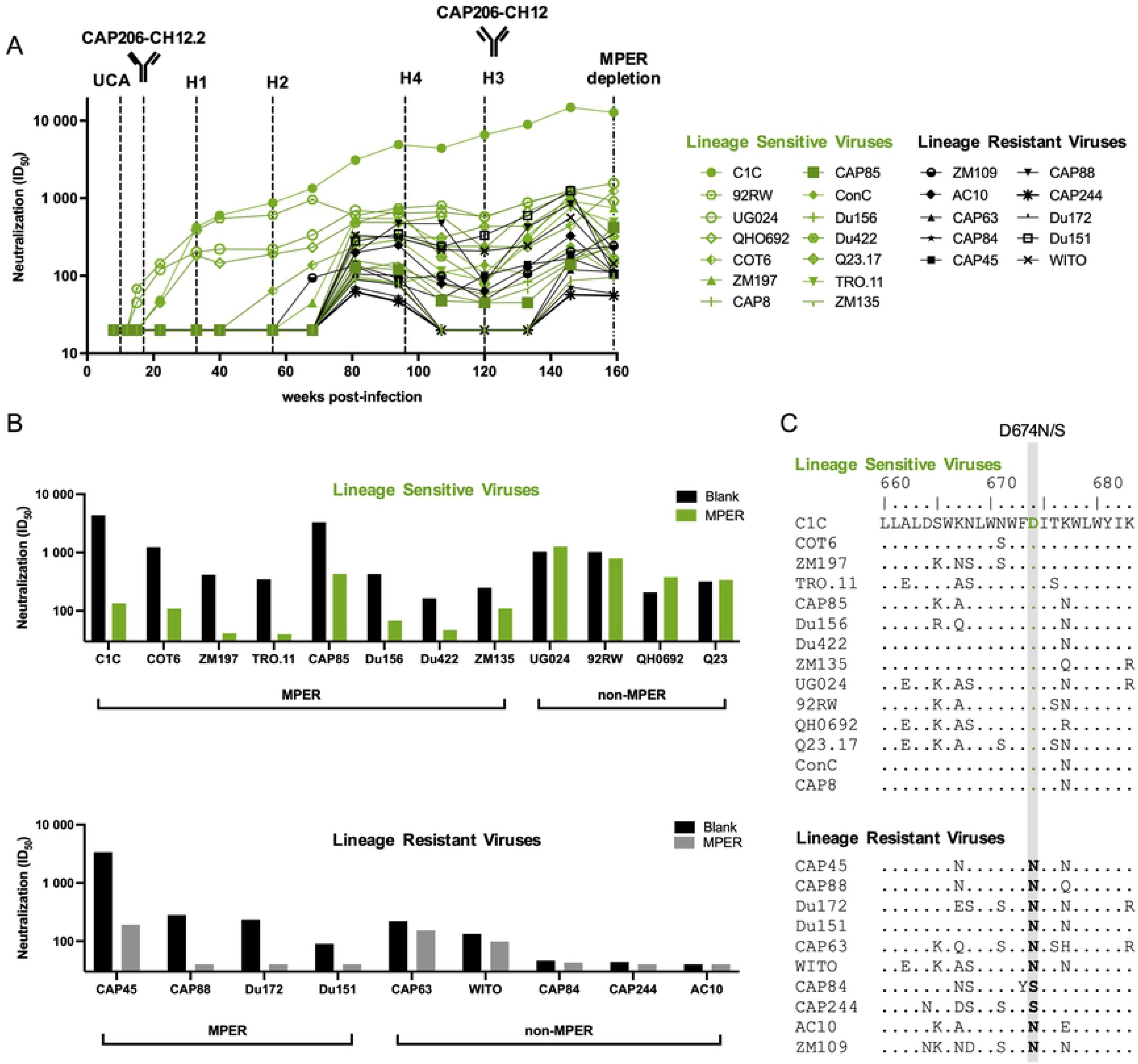
Multiple MPER lineages and specificities in CAP206 plasma. **(A)** Longitudinal plasma sampled over three years (shown as weeks post-infection) were assayed against a multi-subtype panel of 23 viruses sensitive to CAP206 plasma shown as ID_50_ [21]. Viruses shown in green are sensitive to the CAP206-CH12 lineage, while those in black are resistant. The time-points at which the different lineage mAbs (UCA, H2 – H4, CAP206-CH12.2, and CAP206-CH12) were detected/isolated are shown by dashed lines. The time-point at which MPER-depletions were conducted (159 wpi) is shown by the dashed/dotted line. **(B)** Depletion of MPER antibodies from CAP206 serum results in the reduction of neutralization for some but not all viruses. Serum from 159 weeks post-infection was depleted of anti-MPER antibodies using peptide coated beads and tested for neutralization against both lineage-sensitive and lineage-resistant viruses. Results are shown as ID_50_ neutralization with blank bead depleted samples in black and MPER bead depleted samples in green for the lineage-sensitive viruses (top panel) and grey for lineage-resistant viruses (bottom panel). Viruses that showed >2-fold change decrease in neutralization between the blank and MPER-depleted samples were considered to be neutralized by MPER-directed antibodies. Viruses with fold change differences <2-fold were considered non-MPER antibodies. **(C)** Alignment of CAP206-CH12 lineage-sensitive (top panel) and lineage-resistant (bottom panel) viruses compared to C1C, a consensus subtype C MPER sequence. Differences at a single position, 674, distinguished lineage-sensitive viruses, all of which contain a D compared from lineage-resistant viruses containing either an N or S, highlighted in grey.

Magnetic beads coated with the MPR.03 peptide were used to adsorb MPER neutralizing antibodies from CAP206 plasma at 159 wpi (~ 3 years post-infection). We tested the ability of the adsorbed and blank (unadsorbed plasma) to neutralize both lineage-sensitive and lineage-resistant viruses (**Fig 4B**). Within the CAP206-CH12 lineage-sensitive viruses (top panel), the majority (7/11) showed reduced neutralization following depletion (green), confirming MPER specificity [22]. However, four of the viruses that are neutralized by mAbs in this lineage (UG024, 92RW, QH0962, and Q23) showed no reduction of neutralization following MPER-depletion, suggesting other, non-MPER, specificities dominate neutralization of these viruses.

Within the CAP206-CH12 lineage-resistant viruses, five (CAP63, WITO, CAP84, CAP244, and AC10) showed no reduction in neutralization following MPER-depletion, but four (CAP45, CAP88, CAP172, and Du151) did show reduced neutralization titres (**Fig 4B**, bottom panel). This data indicates that at least two antibody specificities, one of which is MPER-directed and the other not, account for the neutralization of lineage-resistant viruses. Overall, this suggests that, in addition to the CAP206-CH12 MPER-directed lineage, donor CAP206 also has another MPER lineage as well as another bNAb with unknown specificity.

An alignment of the MPER sequences between the CAP206-CH12 lineage-sensitive (top panel) and lineage-resistant (bottom) viruses highlights a single amino acid at position 674 (highlighted in grey) that distinguishes these two groups (**Fig 4C**). Where lineage-sensitive viruses have a negatively charged D^674^ compared to the uncharged S^674^ or N^674^ seen in the lineage-resistant viruses. This confirms [22] that the CAP206-CH12 lineage is dependent on D^674^ while the additional MPER lineage within donor CAP206 is not.

### CAP206-CH12 lineage is dependent on a highly variable MPER residue

In order to test the dependence of CAP206-CH12 on D^674^, we introduced mutations at this and other sites within the MPER known to affect sensitivity to CAP206-CH12, VRC42.01, and 4E10 [15,22,25] into the COT6 virus which is sensitive to all three antibodies (**Fig 5A**). We compared the ability of these antibodies (including CAP206-CH12-GW) to neutralize the different mutant viruses compared to the wild-type (COT6).

**Fig 5.**
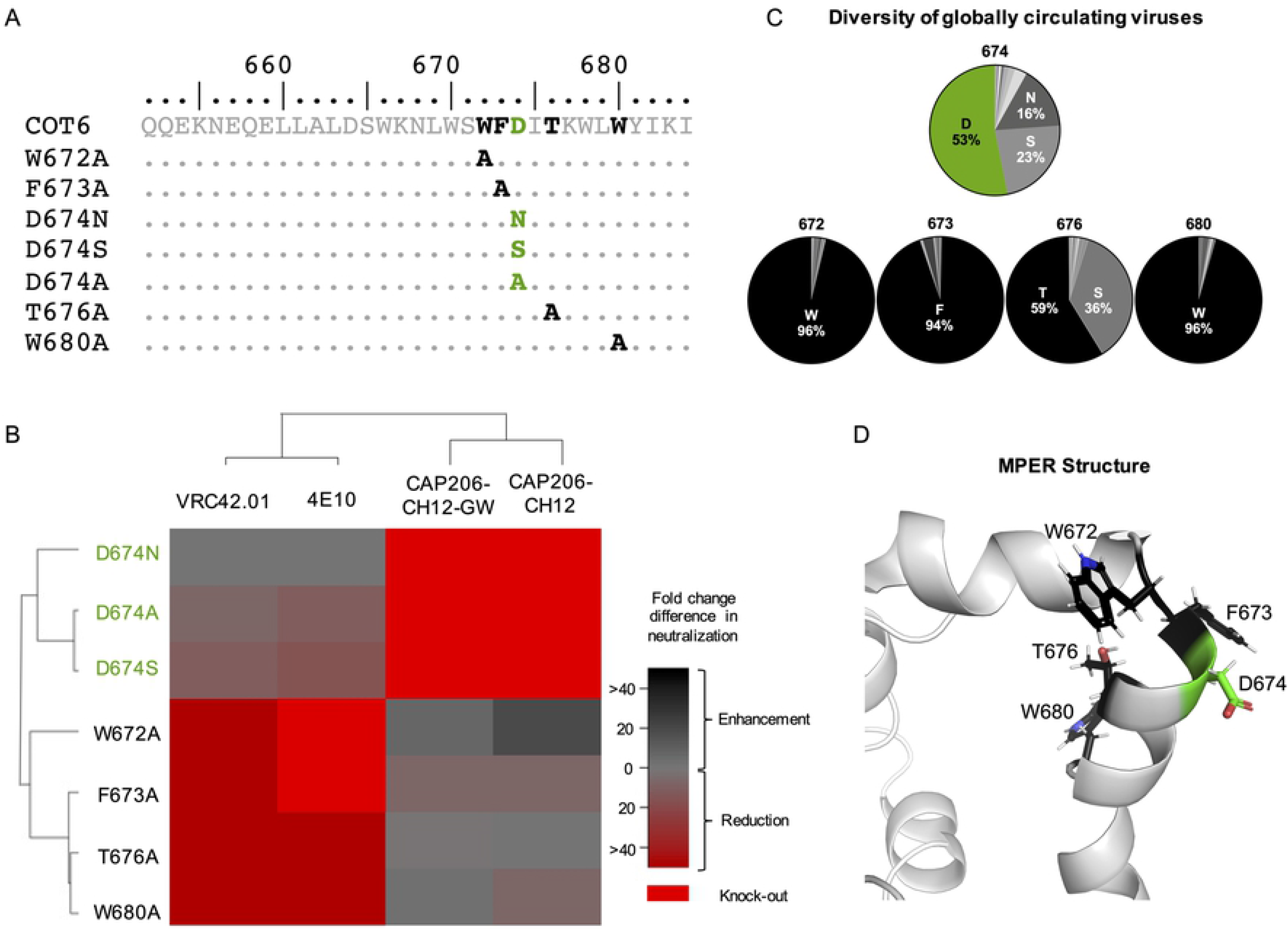
CAP206-CH12 lineage is dependent on the highly variable D^674^ residue. **(A)** MPER sequence alignment of mutations introduced into the MPER of the COT6 virus (sensitive to CAP206-CH12, VRC42.01, and 4E10). Highlighted in black are key amino acid residues for VRC42.01, 4E10 and in green are key residues for CAP206-CH12 neutralization. **(B)** Heat map showing the fold change differences in neutralization between COT6 wild-type and mutant viruses (represented by the rows) for VRC42.01, 4E10, CAP206-CH12, and CAP206-CH12-GW. Red represents reduction or loss of neutralization against the mutant, and black represents increased neutralization against the mutant. **(C)** Pie charts representing the relative frequency of amino acid residues at positions 674 (CAP206-CH12 dependent residue shown in green), 672, 673, 676, and 680 (VRC42.01 and 4E10 dependent residues are shown in black) based on 7,510 MPER sequences, representing all HIV group M subtype sequences from the Los Alamos Database (https://www.hiv.lanl.gov). **(D)** Schematic representation of the MPER “turn” and “kink” structure [25] with key residues for the CAP206-CH12 lineage shown in green and those for VRC42.01 and 4E10 shown in black.

Strikingly, CAP206-CH12 had distinct epitope dependencies (highlighted in green) compared to VRC42.01 and 4E10 (highlighted in black) (**Fig 5A** and **B)**. Both CAP206-CH12 and CAP206-CH12-GW were highly dependent on the negatively charged aspartic acid (D) at position 674, with mutation of this residue to either an A^674^, S^674^, or N^674^ resulting in complete knock-out of neutralization. VRC42.01 and 4E10 showed some reduction (6 – 15-fold) in neutralization when D^674^ was mutated to either an S^674^ or A^674^ but a mutation to an N^674^ had no effect on neutralization. In contrast, VRC42.01 and 4E10 showed a more than 20-fold reduction or complete loss of neutralization when W^672^, F^673^, T^676^, or W^680^ were mutated, whereas these mutations resulted in modest effects on CAP206-CH12 and CAP206-CH12-GW.

When comparing the frequencies of these residues among 7,510 MPER sequences, including all HIV group M subtype sequences, from the Los Alamos Database (https://www.hiv.lanl.gov), we noted that residues 672, 673, and 680, upon which VRC42.01 and 4E10 are dependent are highly conserved residues. In contrast, amino acid 674 upon which CAP206-CH12 and CAP206-CH12-GW show dependence is highly variable, with only 53% of global viruses containing an aspartic acid (**Fig 5A** and **C)**. Furthermore, modeling of the residues which form part of the two MPER sub-epitopes recognized by 4E10/VRC42.01 versus CAP206-CH12 suggests that these antibodies likely target different aspects of the MPER. Although CAP206-CH12, VRC42.01, and 4E10 all target the MPER within the turn of the C-terminal helix [15,22,26], VRC42.01 and 4E10 target residues facing the hydrophobic core whereas CAP206-CH12 targets residues on the outer solvent-exposed surface of the helix (**Fig 5D**). Therefore, CAP206-CH12, VRC42.01, and 4E10 depend on different amino acids, and have different modes of binding to the MPER, likely accounting for the observed differences in breadth and potency.

## DISCUSSION

Antibodies CAP206-CH12, VRC42.01, and 4E10 all target the C-terminal helix of MPER of gp41 and use the same variable germline genes in both the heavy and light chains (*IGHV1-69* and *IGKV3-20*) [15,22,25]. Despite these shared features, VRC42.01 and 4E10 are amongst the broadest antibodies (>95%), whereas CAP206-CH12 only displays ~6% breadth [15]. Here we show, using antibody lineage sequencing, that the CAP206-CH12 lineage undergoes some convergent evolution with VRC42.01 and 4E10, introducing a ^111.2^GW^111.3^ motif within the CDRH3 region of the heavy chain. As with VRC42.01 and 4E10 [15], this ^111.2^GW^111.3^ motif improves neutralization potency within the CAP206-CH12 lineage, but not to the same levels as seen in VRC42.01 and 4E10. We explored the mechanism for limited breadth, and showed that CAP206-CH12 is dependent on a highly variable residue on the solvent-exposed surface on the elbow of the MPER whereas VRC42.01 and 4E10 are dependent on multiple conserved residues facing the hydrophobic core of the MPER. Thus, despite using the same germline genes and demonstrating convergent evolution, the dependence of these lineages on different residues within the same epitope and orientation of binding to the MPER likely determine neutralization breadth and potency.

The CAP206-CH12 lineage contains a string of hydrophobic tyrosine residues next to the ^111.2^GW^111.3^ motif which is encoded by *IGHJ6*. In contrast VRC42.01, and 4E10 use *IGHJ4* which is much shorter and doesn’t contain the string of hydrophobic tyrosine residues. The hydrophobic residues in the CAP206-CH12 lineage may cause a steric clash between the antibody heavy chain and the hydrophobic core of the MPER, perhaps driving this lineage to favor the external solvent exposed region. Thus, the pairing of the *IGHV1-69* with *IGHJ6* instead of *IGHJ4* may have resulted in the CAP206-CH12 lineage initially engaging the MPER in a different orientation than that of VRC42.01 and 4E10 and thereafter maturation dependence on D^674^, leading to narrower neutralization.

We show that donor CAP206 contained multiple bNAb specificities. In addition to the CAP206-CH12 lineage, we have provided evidence for an additional MPER lineage as well as another bNAb specificity to an undefined epitope. We and others have previously described the development of multiple bNAb specificities within a single donor, which contribute towards the overall breadth of these donors [15,27–36]. An effective vaccine would likely need to induce responses to multiple epitopes [2,37] Therefore, studying donors that develop multiple bNAb lineages during infection may provide the insight needed for immunogen design for such strategies.

Neutralizing antibody responses to the MPER tend to be rarer than those targeting other epitopes on the HIV envelope [38]. It is therefore intriguing that both CAP206-CH12 and VRC42.01 were identified within donors that had multiple lineages targeting the MPER [15]. This suggests that there may be certain properties within the viral variants that infected the donors CAP206 and RV217.40512 (the donor from whom the VRC42 lineage was isolated) that gave rise to multiple MPER responses. Due to the highly conserved nature of the MPER, bNAbs targeting this region tend to be among the broadest antibodies isolated to date [39]. Sequential immunogens designed based on variants identified in CAP206 and/or RV217.40512 may enable the development of such responses in a vaccine setting.

Overall, this data helps to explain why the CAP206-CH12 lineage, despite convergent evolution of variable heavy and light chain genes and similar epitope targeting, fails to develop the same level of breadth and potency as VRC42.01 and 4E10. Thus, while shared germline genes and convergent evolution within bNAb classes are crucial aspects of germline-targeting, they may not be sufficient to drive neutralization breadth.

## MATERIALS AND METHODS

### Ethics Statement

CAP206 is a participant enrolled in the CAPRISA 002 Acute Infection study, established in 2004 in Kwa-Zulu Natal, South Africa. The CAPRISA 002 Acute Infection study was reviewed and approved by the research ethics committees of the University of KwaZulu-Natal (E013/ 04), the University of Cape Town (025/2004) and the University of the Witwatersrand (MM040202). CAP206, an adult, provided written informed consent.

### Human samples, viruses, and mAbs

CAP206, a CAPRISA Acute Infection cohort participant, infected with an HIV subtype C virus, developed plasma neutralization breadth [40]. Stored peripheral blood mononuclear cells (PBMCs) taken at 10, 33, 56, 94, 120, 185 and 198 weeks post-infection (wpi) from donor CAP206 were used for CAP206-CH12 lineage tracing using Illumina MiSeq next generation sequencing. Stored serum samples from 159 weeks post-infection (3 years) were used for MPER adsorptions. Stored plasma samples from 8, 12, 15, 22, 33, 40, 56, 68, 81, 94, 107, 120, 133, 146, 159, 185 and 198 wpi were used for longitudinal heterologous neutralization assays. The CAP206-CH12 and CAP206-CH12.2 mAbs [22,23] were obtained as plasmids from Drs Larry Liao and Barton Haynes of Duke University. VRC42.01 and 4E10 were synthesized by GenScript (Hong Kong, China) and subcloned into a SEK vector. Envelope viral clones used in this study were obtained from the NIH AIDS Research and Reference Reagent Program or the Programme EVA Centre for AIDS Reagents, NIBSC, UK. The C1C MPER chimera was obtained from George Shaw (University of Alabama, Birmingham).

### CAP206-CH12 lineage next-generation sequencing

Total RNA was extracted from cryopreserved PBMCs at 7 different time-points (10, 33, 56, 94, 120, 185, and 198 wpi) using the AllPrep DNA/RNA mini kit (Qiagen) as per the manufacturer’s specifications. Reverse transcription was carried out using Random Hexamers (Integrated DNA Technologies) and Superscript III RT enzyme (Invitrogen). Primers specific to CAP206-CH12 heavy and light chains (*IGHV1-69* and *IGKV3-20*) were designed such that forward primers bound in the leader sequence (*IGHV1-69*: 5’- CAGGTSCAGCTGGTGCARTCTGGG-3’ and *IGKV3-20*: 5’-GAAATTGTGTTGACRCAGTCTCCA-3’). Reverse primers bound to CH1 of the constant region (IgG: 5’AGGGYGCCAGGGGGAAGAC-3’, Kappa: 5’- GGGAAGATGAAGACAGATGGT-3’). All primers included Illumina MiSeq barcodes to allow sequencing on the MiSeq. Samples from each time point were amplified seven times for both the heavy and light chains to ensure adequate coverage and minimize PCR bias. PCR conditions were as previously described [19]. Nextera XT unique dual indexing combinations selected from Illumina Indexing Kit V2 Set B were added to the pooled MiSeq amplicon libraries. All products were checked on an Agilent bioanalyzer High Sensitivity DNA kit (Diagnostech) and Qubit dsDNA HS assay (ThermoFisher Scientific) and cleaned-up using 0.75X Ampure Beads (Beckman-Coulter). A final concentration of 4.5 pM denatured DNA library with 10% PhiX control (Illumina) was run on the Illumina MiSeq, using the MiSeq reagent kit (version 3) with 2 x 300 paired-end reads.

### CAP206-CH12 lineage analysis

Paired-end MiSeq reads were merged into full-length reads for each time point using PEAR [41]. Paired sequences were then de-replicated using USEARCH [42] resulting in unique reads. The SONAR bioinformatics pipeline [43] was used to identify clonally related reads to CAP206-CH12. In brief, germline V and J genes were assigned, the CDR3 regions were identified, and identity to CAP206-CH12 and level of somatic hypermutation for each of the reads was calculated. Clonally related sequences were selected based on germline gene usage and CDR3 identity ≥80% to CAP206-CH12. To account for sequencing and/or PCR error, all singletons were removed and the remaining sequences were clustered at 99% identity using USEARCH. A representative from clusters with a size of more than 1% of the most abundant read per time-point was selected for downstream analyses. Tools from the immcantation portal were used to reconstruct the antibody lineage and create the phylogenetic tree [44]. Briefly, clonally related heavy chain sequences were submitted to IMGT High V-quest [45,46] for analysis, and the resulting summary files were then parsed into a Change-O database. The UCA of the lineage was used as the germline-defined sequence for the reconstruction. Within R, the Alakazam package [44] was used to read in the Change-O database and reconstruct the lineage creating clones and introducing inferred sequences using buildPhylipLineage. The resulting tree was plotted using igraph [47]. The edge lengths of the tree were further modified to be proportional to the number of mutations between nodes and all nodes with no mutations from the parent node were deleted using Inkscape v 0.91.

### CAP206-CH12 lineage antibody production

Selected clonally related heavy sequences were ordered codon-optimized from GenScript (Hong Kong, China) and sub-cloned in the CMV/R expression vector [48,49]. All heavy chain clonally related sequences (H1 - H6), were paired with the mature CAP206-CH12 light chain (CH12 L). The UCA heavy chain was paired with the UCA light chain (UCA L). CAP206-CH12 was also paired with two additional light chains (K1 and K2). These heavy and light chain pairs were sub-cloned into an IgG1 backbone, co-transfected, and expressed in HEK293F cells (obtained from the NIH AIDS Research and Reference Reagent Program, Division of AIDS, NIAID, NIH) grown in FreeStyle 293 Expression Medium (Gibco, Thermo Scientific, MA, USA) at 37°C, 5% CO2, 70% humidity and 125 rpm. Cultures were harvested after seven days by centrifugation at 4000 x g, and supernatants were filtered through 0.22 μm before purification by protein A chromatography.

### Site-directed mutagenesis

Specific amino acid changes on envelope clones and CAP206-CH12 heavy chain CDRH3 were introduced using the QuikChange Lightning Site-Directed Mutagenesis Kit (catalog #210519, Agilent Technologies, CA, USA). Mutations were confirmed by sequence analysis.

### Neutralization assays

Neutralization was measured as a reduction in luciferase reporter gene expression after a single round of infection of TZM-bl cells with Env-pseudotyped viruses as previously described [50]. Briefly, monoclonal antibodies (mAbs) or heat-inactivated plasma/serum were incubated with pseudovirus for 1 hour, before the addition of cells for infectivity. Neutralization titres were calculated as the inhibitory antibody concentration (IC_50_) or reciprocal plasma/serum dilution (ID_50_) causing 50% reduction of relative luminescence units (RLU) with respect to the virus control wells (untreated virus). Starting concentrations of mAbs were at 200 μg/mL in order to determine a precise titre.

### Anti-MPER activity

Streptavidin-coated magnetic beads (Dynal MyOne Streptavidin C1; Invitrogen) were incubated with the biotinylated peptide MPR.03 (KKKNEQELLELDKWASLWNWFDITNW LWYIRKKK-biotin-NH2) (NMI, Reutlingen, Germany) at a ratio of 1 mg of beads per 20 μg peptide at room temperature for 30 min. Sera were diluted 1:20 in Dulbecco’s modified Eagle’s medium (DMEM)-10% fetal bovine serum and incubated with the coated beads for one hour at a ratio of 2.5 mg of coated beads per mL of diluted plasma. This was followed by a second adsorption at a ratio of 1.25 mg of coated beads per mL of diluted sample. After each adsorption, the beads were removed with a magnet, followed by centrifugation, and were stored at 4°C. The antibodies bound to the beads were eluted by incubation with 100 mM glycine-HCl elution buffer (pH 2.7) for 30s with shaking and then pelleted by centrifugation and held in place with a magnet. The separated immunoglobulin G (IgG) was removed and placed into a separate tube, where the pH was adjusted to between 7.0 and 7.4 with 1 M Tris (pH 9.0) buffer. The same beads were acid eluted twice more. The pooled eluates were then diluted in DMEM, washed over a 10-kDa Centricon plus filter, and resuspended in DMEM. The adsorbed sera were then used in neutralization assays. The reduction (>32-fold) in neutralization by the MPER-adsorbed plasma (green, top panel **Fig 3B**) compared to unabsorbed plasma (black) against C1C, an HIV-2/HIV-1 MPER chimera involving an HIV-2 backbone (7312A) with an HIV-1 MPER region, demonstrated the efficient removal of anti-MPER antibodies.

### Data Availability

Unprocessed MiSeq antibody sequencing data for the CAP206-CH12 lineage, have been deposited for public access into the National Center for Biotechnology Information (NCBI) Sequence Read Archive (SRA), under the following BioProject accession number: PRJNA757277. The heavy and light chain antibody sequences used to generate the UCA, CAP206-CH12.2, CAP206-CH12 as well as heavy chains of H1 – H6 have been deposited into GenBank, accession numbers: MZ825048 - MZ825058. Viral envelope sequences of heterologous viruses are also available in GenBank, accession numbers: AC10.0.29 - AY835446, CAP8 - EF203976, CAP45 - DQ435682.1, CAP63 - EF203973, CAP84 - EF203963, CAP85 - EF203985, CAP88 - EF203972, CAP244 - EF203978, ConC - DQ401075, COT6 - DQ447266.1, Du151 - DQ411851.1, Du156 - DQ411852, Du172 - DQ411853, Du422 - DQ411854, Q23.17 - AF004885, QH0692 - AY835439.1, TRO.11 - AY835445, UG024 - AAB04896, WITO - AY835451, ZM109 - AY424163, ZM135 - AY424079, ZM197 - DQ388515, 92RW – U88823.1.

## ACKNOWLEDGEMENTS

We thank the donor CAP206 from the CAPRISA Acute Infection cohort and the clinical and laboratory staff at CAPRISA for providing specimens. C.S was supported by the National Institute of Allergy and Infectious Diseases of the National Institutes of Health under Award Number U01AI136677. P.L.M. is supported by the South African Research Chairs Initiative of the Department of Science and Innovation (DSI) and the National Research Foundation (NRF) (Grant No 98341). This work was funded by the Center for HIV/AIDS Vaccine Immunology (CHAVI) (grant A1067854), the South African HIV/AIDS Research and Innovation Platform (SHARP) of the Department of Science and Technology (DST), the Poliomyelitis Research Foundation (PRF), South African Medical Research Council (SAMRC) SHIP program and CAPRISA. CAPRISA was supported by the National Institute of Allergy and Infectious Diseases, National Institutes of Health, U.S. Department of Health and Human Services (grant U19 AI51794).

